# Transcriptional pulsing of a nucleolar transgene

**DOI:** 10.1101/040956

**Authors:** Viola Vaňková Hausnerová, Pavel Křížek, Guy M. Hagen, Christian Lanctôt

**Affiliations:** Institute of Cellular Biology and Pathology, First Faculty of Medicine, Charles University in Prague

**Keywords:** Transcription, nucleolus, pulsing, RNA polymerase II, living cells

## Abstract

The pulsatile nature of transcription has recently emerged as an important property of gene expression. Here we report on the characterization of a RNA polymerase II transgene that is transcribed in the nucleolus. Using the MS2-GFP reporter system and live cell imaging, we found that the synthesis of a MS2-tagged transcript in the nucleolus was discontinuous in all of the cells that were observed, with periods of activity lasting from 15 minutes to 21 hours. The frequency of pulse lengths could be fitted with an exponential function, from which we determined that transcription occurs on average for periods of 20 minutes. These ON periods alternate with periods of inactivity which last on average 29 minutes. The post-mitotic reactivation of transcription was found to be asynchronous in daughter cell pairs. Our observation of discontinuous transcriptional activity in the nucleolus may reflect cycling in the assembly and disassembly of active chromatin structure in and/or around the rDNA genes.

## Introduction

With the recent development of live cell imaging techniques, it has been shown that transcription of individual genes occurs in bursts at the single cell level. This phenomenon has been observed in a range of organisms including bacteria (Golding et al., 2005), yeast (Karpova et al., 2008; Zenklusen et al., 2008), *Dictyostelium* (Chubb et al., 2006; Corrigan and Chubb, 2014), *Drosophila* (Paré et al., 2009), *Arabidopsis* (Moreno-Risueno et al., 2010) and mammals (Raj et al., 2006; Harper et al., 2011; Suter et al., 2011; Larson et al., 2013). Elegant mathematical modeling suggests that the pulsatile nature of gene transcription is linked to intrinsically random events in the gene regulation process (Elowitz et al., 2002; Raj et al., 2006). The chromatin environment also seems to play an essential role in the control of transcriptional pulsing since manipulations of chromatin modifications by histone deacetylase inhibitors led to substantial changes in bursting dynamics (Harper et al., 2011; Suter et al., 2011). The overall transcriptional output of a gene can thus be modulated either by the size of individual bursts (Raj et al., 2006) or by the frequency of bursting (Larson et al., 2013).

Most of the studies on transcriptional pulsing have been concerned with RNA polymerase II (RNA pol II)-driven transcription in the nucleoplasm. To our knowledge, the *in vivo* dynamics of transcription occurring in the nucleolus have not been investigated. The nucleolus is the site of transcription of the ribosomal DNA (rDNA) genes (Perry, 1962; Sirri et al., 2008). The molecular mechanisms involved in the regulation of rRNA synthesis have been extensively investigated using biochemical approaches [reviewed in (Schneider, 2012; Grummt and Langst, 2013)]. Approximately 50% of the rDNA genes are in an open, transcriptionally active state; the other half is tightly occupied by nucleosomes and transcriptionally silenced (Conconi et al., 1989; Grummt and Pikaard, 2003). The promoters of active rDNA genes are hypomethylated and are bound by acetylated histones. This epigenetic signature is established and maintained through binding of a molecular complex that comprises the Cockayne Syndrome Protein B ATPase and the G9a histone methyltransferase (Yuan et al., 2007). Conversely, the promoters of inactive rDNA genes are hypermethylated and bound by methylated histones. The NoRC chromatin remodeling complex plays a crucial role in the establishment of these silencing modifications (Santoro et al., 2002).

Transcription can be imaged in living cells using the bacteriophage-derived MS2-GFP reporter system (Bertrand et al., 1998). This reporter relies on the insertion of multiple 19-nt MS2 stem loops into an RNA of interest and on the expression of a fluorescent fusion protein that binds these repeats with high affinity. This approach has been used to show, among other things, that in mammalian cells transcription from heterologous promoters such as the CMV promoter does not occur in bursts whereas transcription from endogenous promoters does (Yunger et al., 2010). We now report on the transcriptional dynamics of a MS2-tagged transgene that is active inside the nucleolus.

## Materials and Methods

### Cell culture and drug treatment

HeLa cells (ATCC no. CCL-2) were cultured in DMEM containing 4.5 g/l of glucose (Life Technologies, Weltham, USA) and supplemented with 10% FBS, penicillin (50 units/ml) and streptomycin (50 μg/ml). For treatment with actinomycin D, cells were grown to 60-70% confluence on uncoated 18 mm x 18 mm glass coverslips and then incubated with actinomycin D (Sigma-Aldrich, St. Louis, USA) at a concentration of 0.025 μg/ml (0.02 μM) for 1 hour.

### Construction of expression vectors

A 2109 bp blunted AseI fragment from pIRESpuro3 (Clontech, Mountainview, USA) was inserted into the 4808 bp blunted ClaI-XhoI fragment of pMS2-GFP (a kind gift from Dr. R. H. Singer, Albert Einstein College of Medicine) in order to generate a vector expressing a bicistronic mRNA encoding MS2-GFP and puromycin acetyltransferase (pMS2GFP-IRES-PURO). A 654 bp BamHI-BglI fragment from pSL-MS2-12X (a kind gift from Dr. R. H. Singer) was cloned into the BglII site of pEGFP-GPI (a kind gift of Dr. G. Kondoh, Kyoto University), after which the EcoRI-XhoI EGFP fragment was replaced with a PCR fragment encoding G418-resistance. The resulting vector (pNEO-MS2^12X^) contains a CMV-chicken β actin (CAG) promoter driving the expression of the neomycin phosphotransferase coding sequence fused at the 3’ end to 12 repeats of the MS2 stem loop.

### Generation of cell line

Three micrograms of the pMS2GFP-IRES-PURO vector were transfected into ∼10^6^ HeLa cells using the FuGene HD reagent according to the manufacturer’s instructions (Roche Applied Science, Basel, Switzerland). Cells were selected by treatment with 2 μg/ml of puromycin (Sigma-Aldrich, St. Louis, USA) for 5 days. The pool of resistant colonies was expanded and ∼8x10^5^ of these cells were transfected with 2 μg of pNEO-MS2^12X^ using FuGene HD as before. G418 selection (500 μg/ml, 74.1% potency, purchased from Life Technologies) was begun 2 days after transfection and lasted 14 days. Individual clones were grown after limiting dilution of the G418-resistant pool. A total of 44 clones were screened; two displayed a prominent MS2-GFP spot in the nucleus and were subsequently referred to as MS2^12x^-12 and MS2^12x^-15. To assess the integrity of the transgene, genomic DNA was extracted with DNeasy Blood and Tissue Kit (Qiagen, Hilden, Germany) and analyzed by PCR using either CloneAmp HiFi PCR Premix (Clontech, Mountain View, USA) or Q5 DNA polymerases (New England Biolabs, Ipswich, USA).

### Immunofluorescence

The mouse monoclonal antibody against Ser5-phosphorylated CTD of human RNA polymerase II (CTD4H8) was purchased from Millipore (Billerica, USA). The human antibody against fibrillarin was an autoimmune serum. Secondary antibodies produced in goat and conjugated to Cy5 (anti-mouse) or Cy3 (anti-human) were purchased from Jackson ImmunoResearch (Baltimore, USA). All incubations were performed at room temperature. HeLa cells grown on uncoated 18 mm x 18 mm coverslips were fixed with 4% formaldehyde in 1X PBS for 10 minutes. Cells were washed with 1X PBS and permeabilized with 0.2% (v/v) Triton X-100 for 10 minutes. After blocking with 1% (w/v) BSA in 1X PBS (PBS-BSA) for 30 minutes, cells were incubated with primary antibodies for 1 hour (1/200 dilution in PBS-BSA). After 3 washes with 1X PBS for 5 minutes, cells were incubated with secondary antibodies for 30 minutes (1/400 dilution in PBS-BSA). After final washes, samples were counterstained with DAPI (1 μg/ml) and mounted in Prolong Gold (Life technologies, Weltham, USA).

### Chromatin immunoprecipitation (ChIP)

ChIP was performed as described previously (Diermeier et al., 2013; Duskova et al., 2014). In brief, cells were fixed with 1% formaldehyde in 1X PBS, lysed in SDS lysis buffer (1% SDS, 50 mM Tris-HCl pH 8.0, 20 mM EDTA, protease inhibitors) and chromatin was sonicated using a Bioruptor sonicator (Diagenode, Denville, USA). The rabbit polyclonal antibody against the RPA194 subunit of human RNA polymerase I and the normal rabbit IgG were purchased from Santa Cruz Biotechnology (Dallas, USA). The mouse monoclonal antibody against the CTD of human RNA polymerase II (8WG16) was purchased from Abcam (Cambridge, UK). Antibodies (5 μg), chromatin and protein A/G agarose beads (Santa Cruz Biotechnology, Dallas, USA) were diluted in IP buffer (20 mM Tris-HCl, 2 mM EDTA, 1% Triton X-100, 150 mM NaCl, pH 8.0, protease inhibitors) and incubated at 4°C overnight. Beads were washed in IP buffer, then in LiCl buffer (0.25 M LiCl, 1% NP40, 1% Deoxycholate, 1 mM EDTA, 10 mM Tris-HCl, pH 8.0), and finally in TE buffer (10 mM Tris-HCl, 1 mM EDTA pH 8.0). Immunocomplexes were eluted using 70 mM Tris-HCl, 1 mM EDTA, 1.5% SDS, pH 6.8, reverse crosslinked by incubation in 200 mM NaCl at 65°C and treated with 100 μg/ml RNAse A and 80 μg/ml proteinase K. DNA was isolated by Qiaquick PCR Purification Kit (Qiagen, Hilden, Germany) and analyzed by qRT-PCR. Results were normalized as percentage of input. The oligos used for ChIP were: for rDNA detection - forward: 5-ACGGTCGAACTTGACTATCTAGAGG-3′, reverse: 5′- CGGAAACCTTGTTACGACTTTTACTT-3′, for Mypn promoter detection - forward: 5′- TGCAGGCTAAGAACATCGGT-3′, reverse: 5-GTGCCATGAAGGAAATGTGA-3′, for MS2^12x^-12 and MS2^12x^-15 transgene detection - forward: 5′-CCCTGGCTCACAAATACCAC-3′, reverse: 5-GAGTCGACCTGCAGACATGG-3′.

### RNA fluorescence in situ hybridization (RNA FISH)

HeLa cells were grown on uncoated 18 mm x 18 mm coverslips and fixed with 4% formaldehyde in 1X PBS for 10 minutes at room temperature. After washing in 1X PBS, samples were permeabilized in 70% ethanol overnight at 4°C. Prior to hybridization, samples were rehydrated in 10% formamide/2X SSC for 30 minutes. The hGAPDH mRNA signal was detected by a Quasar 570-labelled Stellaris RNA FISH probe (Biosearch Technologies, Petaluma, USA). The probe against the MS2 stem loop is a Cy5-labeled oligonucleotide (5-ACATGGGTGATCCTCATGT-3). The hybridization mixture contained the probes against MS2 and hGAPDH (at final concentrations of 0.157 μM and 0.125 μM, respectively), 20 mM ribonucleoside vanadyl complex (New England Biolabs, Ipswich, USA), 2X SSC, 10% dextran sulfate and 10% formamide. Hybridization was carried out for 6 hours at 37°C. After washing with 10% formamide/2X SSC for 30 minutes at 37°C and with 2X SSC for 5 minutes at room temperature, samples were stained with DAPI (1 μg/ml) and mounted in Prolong Gold.

### Image acquisition and live cell fluorescence microscopy

For immunofluorescence and RNA FISH, images were acquired on an Olympus IX71 inverted epifluorescence microscope using a 60X/1.35NA oil immersion objective, an Andor Clara CCD camera (Andor Technology, Belfast, UK) and an automated piezo-Z stage (Prior Scientific Instruments, Cambridge, UK). Live cell imaging was performed using an Andor Revolution system mounted on an Olympus IX81 microscope equipped with a Yokogawa CSU-X spinning disk confocal unit, an iXon Ultra Andor EM-CCD Camera, an automated XY stage and a piezo-Z stage from Prior Scientific Instruments. Images were acquired using a UPLSAPO 60X/1.3 NA silicon oil immersion objective. Excitation light at 488 nm was from a 50 mW solid state laser. Fluorescence was collected using a 525/50 nm bandpass filter. For acquisition of time lapse movies, cells were seeded on untreated MatTek glass bottom dishes (MatTek Corporation, Ashland, USA) and grown in medium without phenol red. *xyz* stacks of ∼114 x 114 x 10 μm^3^ were acquired at multiple positions every 15 minutes for 23-24 hours. All images were acquired at 37°C and in a 5% CO2 atmosphere using a live cell observation chamber (Okolab, Naples, Italy).

### FRAP analysis

FRAP was performed using the FRAPPA module on our Andor Revolution system. Images were acquired using an Olympus UPLSAPO 100X/1.4NA oil immersion objective. A region of interest (ROI, a square of approximately 1.5 x 1.5 μm^2^) containing the MS2-GFP spot was bleached using a 405 nm diode laser (100 mW). Prebleach and postbleach images were acquired for 2 and 12.5 minutes, respectively. In both cases, *xyz* stacks of about 68 x 68 x 3 μm^3^ centered on the MS2-GFP spot were captured every 5 seconds. Data was corrected for overall nucleoplasmic signal using the ImageJ FRAP profiler plugin developed by the Hardin Lab (http://worms.zoology.wisc.edu/research/4d/4d.html#frap). A smaller square ROI was drawn around the transcription spot so that the signal which reappeared after the bleaching and which may have moved laterally was within this ROI. A bigger ROI was drawn around the whole nucleus and was used to define the nucleoplasmic signal for the FRAP profiler plugin.

### Image analysis

We processed images from RNA FISH using ImageJ (http://imagej.nih.gov/ij/). The percentage of cells containing any number of hGAPDH or MS2 transcription spots was determined visually and the DAPI counterstain was used to ascertain their nuclear localization. Images were acquired and processed identically for both untreated and actinomycin D-treated samples.

Time series images were analyzed using custom designed software written in MATLAB. Data processing and measurements were performed on maximum intensity Z-projection images. To segment nuclei, we applied a Gaussian filter followed by an algorithm based on adaptive threshold selection. Once the nuclei had been segmented, spot-like signals were localized by finding local intensity maxima in images that had been previously smoothed with a Gaussian filter of full width at half maximum (FWHM) equal to a user-defined average spot size. Spots were further processed only if their peak signal-to-noise ratio (PSNR) was above a user specified threshold (usually set at 5). At this stage, the PSNR was estimated according to PSNR = (*I*_max_ - μ) / *ρ*, where *I*_max_ is the maximum intensity in the localized spot, *μ* is the average intensity of the nucleoplasmic signal for a given nucleus, and *ρ* is the level of the nucleoplasmic background computed as the standard deviation of all intensity values within a given nucleus. The intensity, size and precise position of the spots were then determined by fitting a symmetric two-dimensional Gaussian function 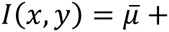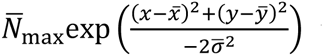 to the raw image data using non-linear least-squares methods (Ovesný et al., 2014). Here *I*(*x,y*) are intensity values of the image at integer coordinates *x,y*, the fitted parameter 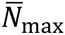 is the maximum spot intensity, 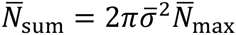 corresponds to the sum of all intensity values in the spot, 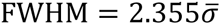 is the fitted size of the spot, coordinates 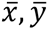 correspond to the fitted center of the spot, and 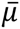 is the average nucleoplasmic intensity in the local neighborhood of the spot. The “spot-to-nucleoplasm” signal strength is given as a signal-to-noise ratio (SNR) according to 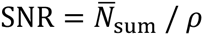. The software generates a time lapse movie of maximum intensity Z-projections in which nuclei are segmented and transcriptional spots are labelled at each time point. These output files were used to visually validate the results of the algorithm.

## Results

### RNA polymerase II-driven transcription of a MS2-tagged transgene that is localized in the nucleolus

As part of a study on the impact of environmental factors on transcriptional pulsing, we generated a vector expressing a neomycin phosphotransferase cassette fused to 12 repeats of the MS2 stem loop in the 3’ untranslated sequence and under the control of the CAG promoter (Fig. 1A) (Niwa et al., 1991). This construct was transfected in a HeLa cell line which stably expresses the MS2 coat protein fused to green fluorescent protein and to a nuclear localization signal (MS2-GFP). After selection and screening, two clones were found that contained a single dot-like MS2-GFP signal in the nucleus. We refer to these clones as MS2^12X^-12 and MS2^12X^-15. Interestingly, not all cells displayed a nuclear dot-like MS2-GFP signal; the proportions of cells that do were on average 41% ± 7% (231 cells analyzed) for MS2^12X^-12 and 33% ± 10% (261 cells analyzed) for MS2^12X^-15. The intensity of the signal was mostly uniform in the population of positive cells. We noticed early on that the dot-like MS2-GFP signal appeared to be localized in the nucleolus in both clones. Immunostaining for fibrillarin, a pre-rRNA processing enzyme, confirmed the presence of the MS2-GFP spot in the nucleolus (Fig. 1B).

**Figure 1.**
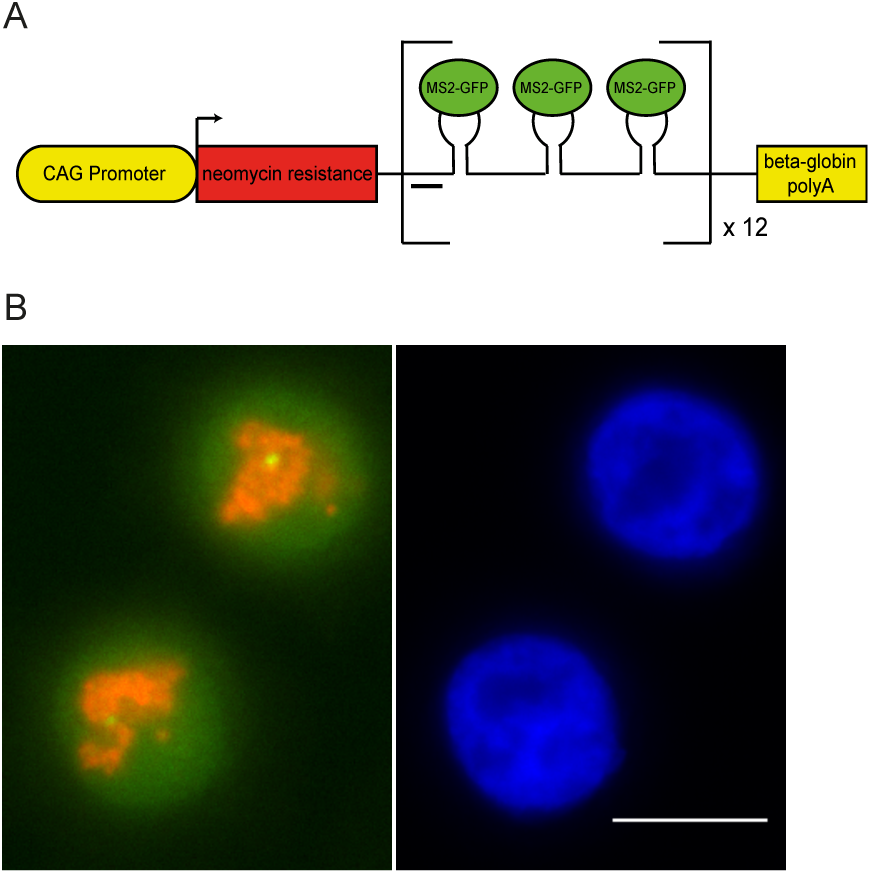
The MS2-tagged transgene is active in the nucleolus. **A)** Scheme of the expression unit. The CMV-chicken β actin (CAG) enhancer and promoter sequences drive the expression of the neomycin phosphotransferase coding region fused to 12 MS2 stem loops in the 3′ untranslated region. **B)** Immunostaining of HeLa MS2^12x^-12 cells with an antibody against fibrillarin, a nucleolar marker. Left panel: MS2-GFP (green) accumulates in a fibrillarin-positive compartment (red). Right panel: DAPI-stained DNA (blue) of the same cells. Single optical sections are shown. Bar, 10 μm.

The unexpected nucleolar localization of the MS2-GFP signal prompted us to ascertain the integrity of the transgene and to determine whether it was indeed being generated by RNA pol II. Analysis of genomic DNA by PCR revealed the presence of the entire expression cassette, with the exception of the 3’ untranslated sequence (not shown). Importantly, the MS2 sequences were found to be linked to the CMV-chicken β actin (CAG) enhancer and promoter sequences. We used chromatin immunoprecipitation (ChIP) to determine which of RNA pol I or RNA pol II was active on the MS2 sequences. In both clones the MS2 sequences could be immunoprecipitated with an antibody against the C-terminal domain (CTD) repeats of RNA polymerase II (Fig. 2A). Immunoprecipitation with an antibody against the large subunit of RNA polymerase I led to the recovery of rDNA sequences, but not of the MS2 repeats. These results strongly suggested that RNA pol II is responsible for the synthesis of the MS2-tagged RNA in the nucleolus. To further confirm these results, we carried out immunofluorescence with an antibody against the initiating form of RNA pol II (phospho-Ser5 on the CTD), reasoning that a signal should be detected at the MS2-GFP dot if RNA pol II was active on the MS2 sequences, as shown by the ChIP data. This was found to be the case (Fig. 2B). A faint but reproducible RNA pol II signal co-localized with the MS2-GFP nucleolar signal in 60 out the 66 cells that were analyzed (91%). In the cells that lack a MS2-GFP dot in the same population, no accumulation of RNA polymerase II was detected in the nucleolus (71 cells analyzed).

**Figure 2.**
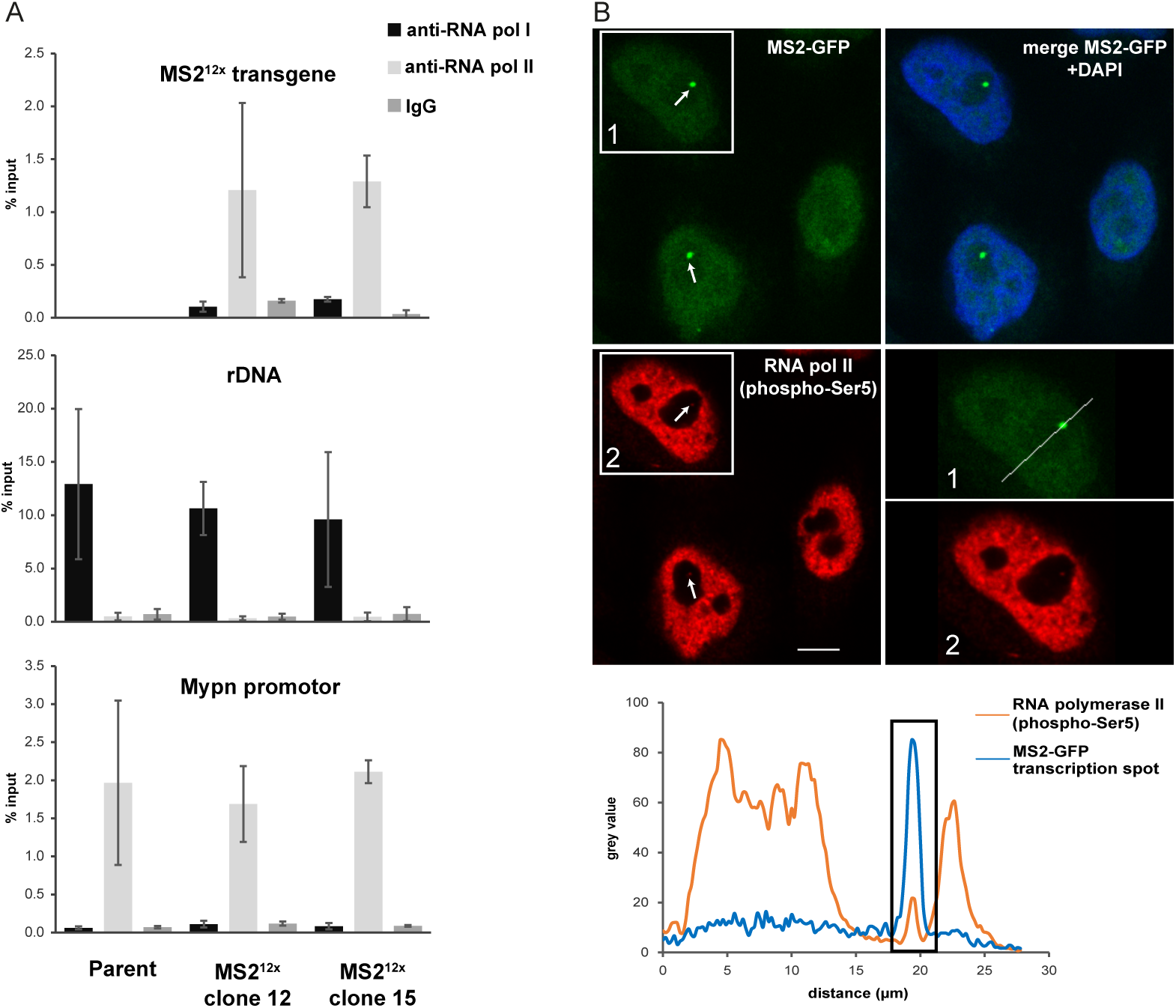
The MS2-tagged nucleolar transgene is transcribed by RNA polymerase II in HeLa MS2^12x^ cells. **A)** Chromatin immunoprecipitation with antibodies against the RPA194 subunit of RNA pol I, the CTD of RNA pol II (clone 8WG16) and control serum (IgG). The qPCR target sequences are indicated above each chart. The MS2-tagged transgene that is active in the nucleolus is bound by RNA pol II (n = 2). Positive controls were performed for RNA pol I binding to rDNA (n = 3) and RNA pol II binding to the promoter of the nucleoplasmic *MYPN* gene (n = 2). **B)** Immunostaining of the HeLa MS2^12x^-15 cells with an antibody against the C terminal domain (CTD) of RNA pol II phosphorylated on Ser5 (bottom left panel, red). The MS2-GFP (top left panel, green) and DAPI + MS2-GFP (top right panel) signals are also shown. Faint accumulation of RNA pol II can be detected at the nucleolar MS2-GFP spot (arrows). Higher magnification of the boxed cell is shown in the bottom right panel. Single optical sections are shown in all cases. Bar, 10 μm. Below are the plot profiles for the MS2-GFP (blue) and RNA pol II (orange) signals across the line drawn in the bottom right panel. The rectangle highlights the colocalization of these two signals.

Having serendipitously generated a cell line which displays RNA polymerase II-mediated transcription in the nucleolus, we decided to study this puzzling cellular activity in more detail. First, we determined whether it is disrupted under conditions that are known to affect nucleolar structure and activity. Cells were incubated in low concentrations of actinomycin D, a treatment which preferentially inhibit nucleolar transcription (Schoefl, 1964; Perry and Kelley, 1970). To assess the extent of transcriptional inhibition, we performed sensitive RNA FISH using fluorescently-labelled probes against the MS2 stem loop sequence or, as a control, against human glyceraldehyde 3-phosphate dehydrogenase (hGAPDH). Treatment with actinomycin D at 0.025 μg/ml for 1 hour led to a more than ten-fold decrease in the proportion of MS2-labeled cells (from 65% to 6% of cells, Fig. 3). The intensity of the signal in the cells that remained MS2-positive after actinomycin D treatment (6% of the population) was greatly decreased (not shown). Neither the proportion of cells displaying hGAPDH transcription sites nor the intensity of these nuclear signals were affected under these conditions. Taken together, these results suggest that RNA pol II-mediated transcription responds to conditions that inhibit RNA pol I when transplanted into the nucleolus.

**Figure 3.**
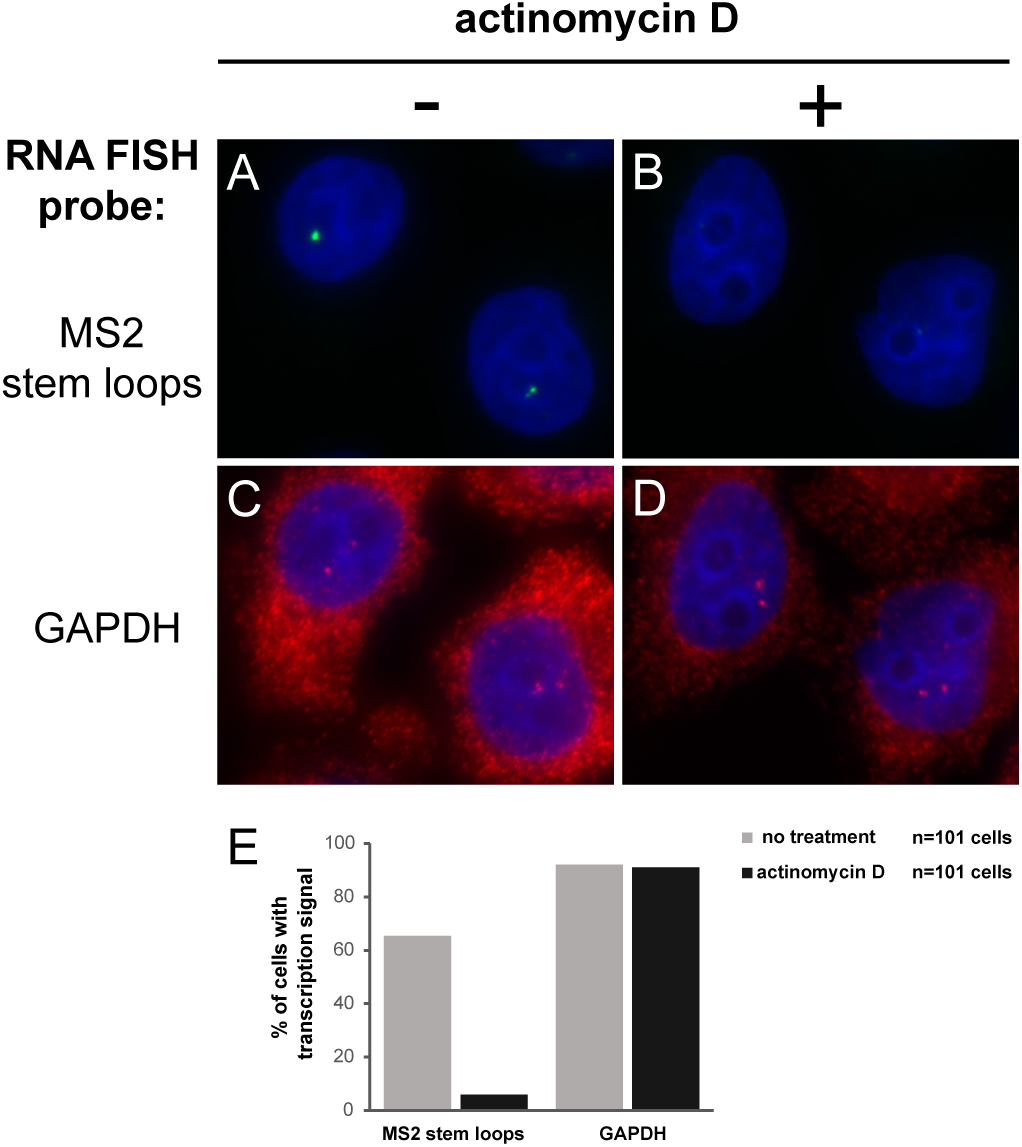
Nucleolar transcription of the MS2-tagged transgene is sensitive to low concentrations of actinomycin D. RNA FISH was performed with probes against the MS2 stem loops (**A-B**, pseudo-colored in green) or hGAPDH (**C-D**, pseudo-colored in red). Cells were treated with 0.025 μg/ml actinomycin D to inhibit nucleolar transcription (**B, D**). DNA was counterstained with DAPI (blue). Bar, 10 μm. Note that the MS2-tagged transcripts are detected only at the transcription site in the nucleus. For hGAPDH, both nascent and cytoplasmic mRNAs are detected. **E**) Proportion of cells with at least one labeled transcription site in the nucleus for each probe and for control and actinomycin D-treated cells (101 cells analyzed).

### Dynamics of the MS2-GFP-labeled transcription spot as measured by FRAP

We used the fluorescence recovery after photobleaching (FRAP) method to measure the dynamics of the MS2-tagged nucleolar transcripts. The MS2-GFP spot was selectively bleached, after which a ∼3 μm thick nuclear cross-section that included the bleached region was scanned on a high-speed spinning disk confocal microscope at 5-second intervals. Signal recovery was detected as early as 2 minutes after bleaching, indicating that active transcription is ongoing at the MS2-GFP spot (Fig. 4A). FRAP curves exhibit a rapid recovery in the first minute followed by a slower recovery phase (Fig. 4B and Fig. S1). The FRAP data is characterized by a large variability, especially at later time points, indicating important differences in the speed and extent of fluorescence recovery between individual cells. Indeed, for a significant fraction of cells the recovery was not complete at the end of the imaging period (12 minutes). On average, it reached a maximum of 76% and 65% of the initial value for clones MS2^12x^-12 and MS2^12x^-15, respectively. This result suggests that up to 35% of the dot-like MS2-GFP signal consists of relatively long-lived RNA molecules that are retained at the site of transcription. Presumably, multiple processes are being imaged simultaneously during the FRAP experiment, i.e., RNA synthesis, processing and degradation. The persistence and/or degradation of the MS2-containing fragments at the site of transcription would prevent their appearance away from the nucleolar spot. This interpretation is consistent with our inability to detect MS2-labelled RNA in the cytoplasm by RNA FISH (see Fig. 3).

**Figure 4.**
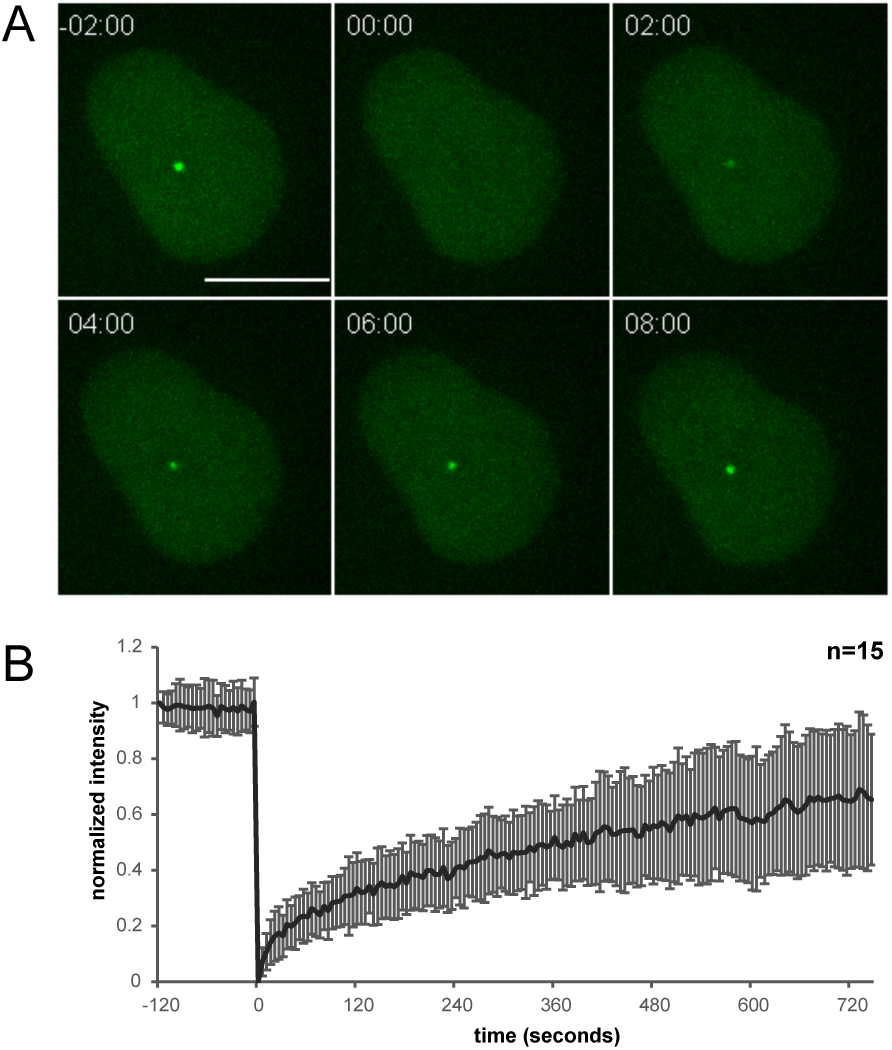
FRAP analysis of the MS2-GFP transcriptional dot. **A)** A representative image series showing reappearance of the MS2-GFP dot-like signal within 2 minutes after bleaching. Shown are maximum intensity projections of image stacks. Time is indicated in minutes and seconds. Bar, 10 μm. **B)** Average intensity profile during FRAP experiments performed on 15 cells of clone MS2^12x^-15. Values (± s.d.) are corrected for total nucleoplasmic signal. Scans were taken every 5 seconds for 14.5 minutes. Maximum intensity projections were used for the evaluation.

### Transcription of the MS2-labeled nucleolar transgene occurs in pulses in living cells

The acquisition of 3D time lapse movies for 24 hours in 15-minute intervals allowed us to analyze the temporal behavior of the MS2-GFP-labelled transcripts. A total of 268 cells were imaged from 21 movies acquired in 3 different experiments. The transcription spots were tracked and quantified using a custom MATLAB-based program (Fig. S2, see Materials and Methods for details). Putative spots were identified as local intensity maxima in the nucleus. Only those with a spot-to-nucleoplasm intensity ratio higher than a user-defined value (not less than 4) were included in our analysis. Signal strength was determined computationally and corresponds to the sum of intensities of all pixels belonging to the Gaussian-fitted spot. Expression was detected in 92% of cells at one point or another. Transcription was discontinuous in all of the cells we observed, with ON periods ranging from less than 15 minutes to 21 hours. The pulsatile nature of transcription is illustrated on Fig. 5A. The interphase imaging data set is represented from the longest to the shortest in Fig. S3. The distributions of the lengths of ON (n = 1104) and OFF (n = 876) periods are plotted on Fig. 5B, C. Fitting these curves to an exponential function leads to an average pulse length of 20.3 minutes ± 1.1 minute and an average OFF time of 29.3 minutes ± 1.7 minute. The relationship between pulse length and the maximum signal strength reached during the pulse is shown in Fig. 5D. The signal strength correlates with pulse length for durations of up to 300 minutes, at which time it reaches a maximum value.

**Figure 5.**
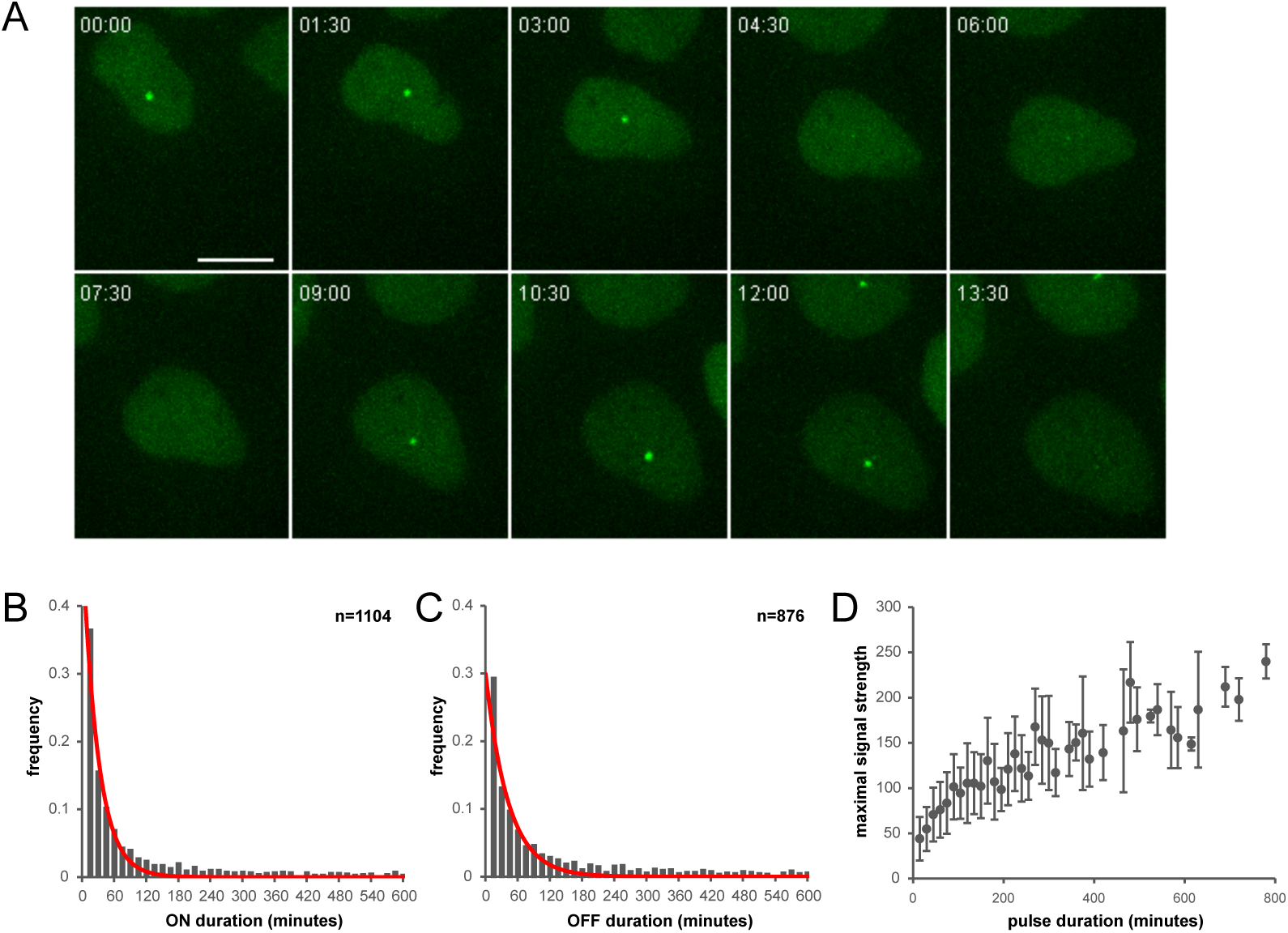
Transcription of the MS2-tagged RNA occurs in pulses. **A)** A representative image series showing the appearance and disappearance of the MS2-GFP transcription spot over time. Shown are maximum intensity projections of image stacks. Time is indicated in hours and minutes. Bar, 10 μm. **B)** Frequency distribution for the duration of ON periods (1104 pulses imaged from 268 cells). The exponential fit of the experimental data is shown in red. **C)** Frequency distribution for the duration of OFF periods (876 pulses from 268 cells). The exponential fit of the experimental data is shown in red. **D)** Plot of the maximum signal strength (relative units) that is reached during pulses of increasing duration. The plotted data is the average ± s.d. A plateau is reached when the pulse duration reaches ∼300 minutes.

We often observed progressive increases and decreases of the MS2-GFP spot intensity during the ON periods. To better characterize this behavior, we analyzed more closely the signal strength for the first and last time points of pulses. These values were divided by the maximum signal strength that is reached at any time during the pulse, and these ratios were used as a rise index (in the case of the first time point of the pulse) and a decline index (in the case of the last time point of the pulse). For short pulses, i.e., for those lasting between 45-60 minutes (n = 187), the rise index was 0.75 ± 0.23 and the decline index was 0.76 ± 0.22 (Fig. 6A, C-D). In contrast, for pulses lasting more than 120 minutes (n = 240), the rise and decline indexes were significantly lower at 0.48 ± 0.22 and 0.52 ± 0.25, respectively (two tailed Student’s *t*-test, p < 0.001, Fig. 6B, C-D). Hence, the shorter the pulse, the more abrupt are the increase and decrease in the strength of the transcription signal.

**Figure 6.**
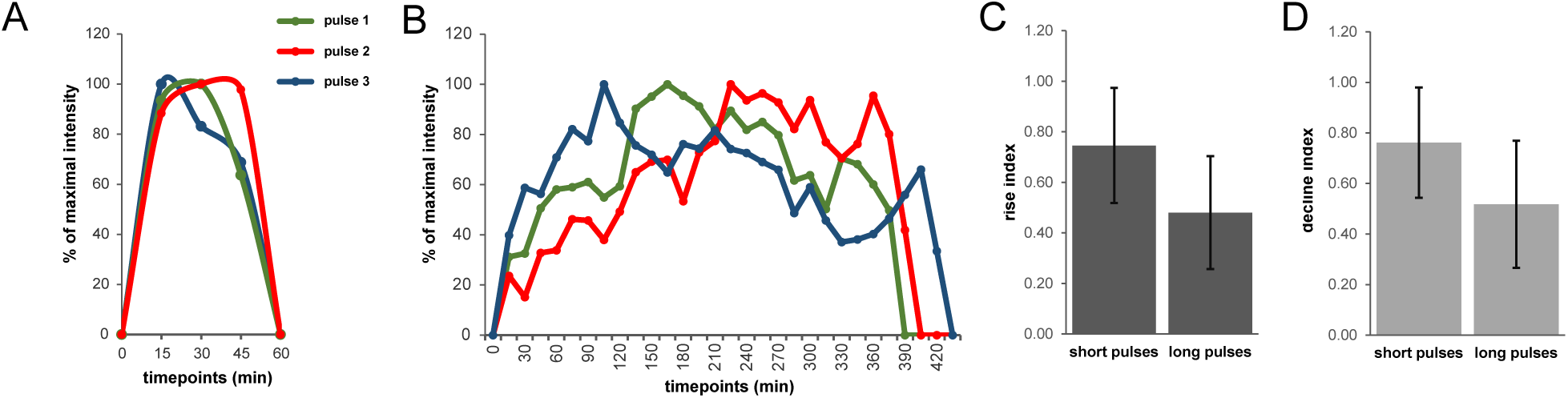
Progressive increase and decrease of transcription during short or long pulses. **A, B)** Normalized intensity profiles for 3 typical short (45-60 minutes) and long (more than 120 minutes) transcriptional pulses, respectively. **C, D)** Average normalized intensities of the first (rise index, dark grey) and last (decline index, light grey) MS2-GFP spot for both classes of transcriptional pulses. Values (± s.d.) are calculated from 187 short pulses and 240 long pulses.

### Transcription resumes asynchronously in daughter cells after cell division

Finally, our dataset allowed us to follow a large number of cells through mitosis and to determine the extent of transcriptional memory from mother to daughter cells (Fig. 7A, B). From a total of 121 daughter cell pairs that were imaged, 16 pairs (13.2%) did not show any expression over the entire imaging period and only 9 (7.4%) resumed transcription of the MS2-tagged RNA synchronously in both daughter cells. The 96 remaining pairs (79.3%) showed asynchronous resumption of transcription after cell division. From the available imaging data, the signal could be detected in only one of the daughter cells for 28 of these pairs. For the rest (n = 68), the MS2-GFP spot could be detected in the two daughter cells, with differences in the onset of expression ranging from 15 minutes to 13.5 hours (Fig. 7C).

**Figure 7.**
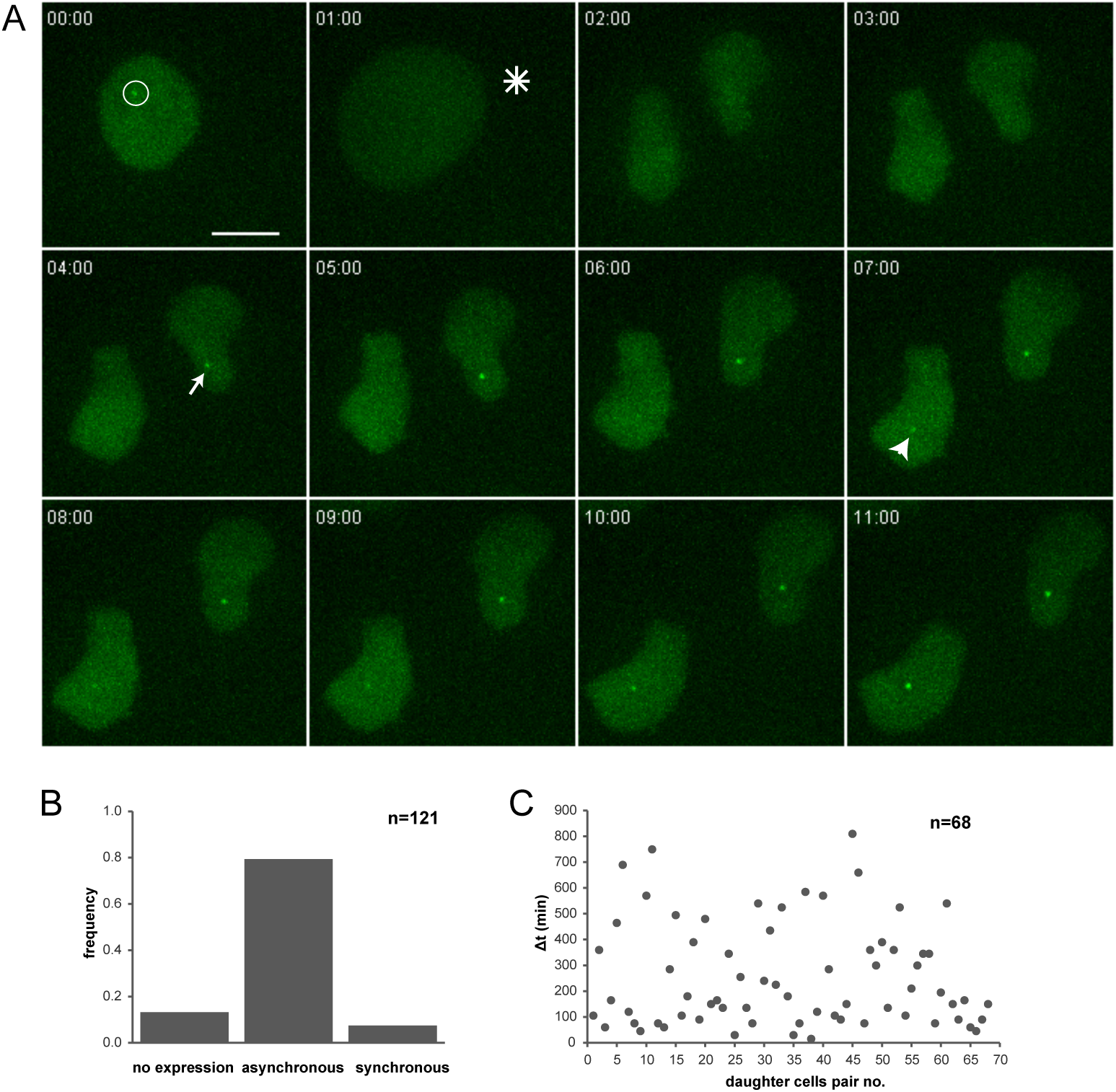
Onset of transcription after mitosis is asynchronous in daughter cells. **A)** A representative image series through mitosis (asterisk) and in interphase. Transcription resumes in one of the daughter cells (arrow) three hours before it does in the other (arrowhead). Shown are maximum intensity projections of image stacks. The transcription spot is circled in the mother cell. Time is indicated in hours and minutes. Bar, 10 μm. **B)** The frequency of each type of event for 121 daughter cell pairs that were imaged, i.e., no expression, asynchronous transcription, synchronous transcription. **C)** Time intervals between the onset of post-mitotic transcription in daughter cells (68 pairs of cells analyzed).

## Discussion

The main result of the present work is the observation of transcriptional pulsing in the living nucleolus. This observation was made possible by the use of the MS2-GFP reporter system, which allowed us to image the transcription of a MS2 tagged transgene stably inserted in the genome of HeLa cells. Since this transgene is under the control of well-characterized enhancer and promoter sequences known to recruit RNA pol II, the fact that its transcription occurs in the nucleolus was an unexpected finding. Nonetheless, the nucleolar RNA pol II activity was confirmed both by ChIP data showing that the MS2 repeats were bound by this polymerase, but not by RNA pol I, as well as by immunostaining results showing the presence of RNA pol II at the nucleolar transcription site of the MS2-tagged transgene. The observation of RNA pol II activity in the cell nucleolus raises a number of intriguing questions. Although it is now generally thought that the nucleolus carries functions other than ribosome biogenesis, such as cell cycle regulation, mRNA processing and viral metabolism (Pederson, 1998; Shou et al., 1999; Hiscox, 2002), few reports have hinted at the possibility of RNA pol II-mediated transcription in the mammalian nucleolus. Especially noteworthy in this regard is the work of Gagnon-Kugler and colleagues, who detected cryptic RNA pol II activity on rDNA in mammalian cells devoid of CpG methylation (Gagnon-Kugler et al., 2009). In yeast, it has been shown some time ago that reporter genes inserted into yeast rDNA repeats can be transcribed by RNA pol II in the absence of RNA pol I activity (Cioci et al., 2003). In fact, in this lower eukaryote a highly conserved sequence in the spacers that separate the individual rDNA units was found to be bidirectionally transcribed by RNA pol II (Kobayashi and Ganley, 2005). Recently, Cesarini and colleagues reported that the transcription levels of this non coding RNA by RNA pol II correlated with the acetylation of Lys-16 of histone H4 on rDNA (Cesarini et al., 2012). It should be noted that the observations of nucleolar RNA pol II activity in yeast were made in mutant cells. The results presented here complement these previous studies by showing that RNA pol II can act in the mammalian nucleolar environment in normal conditions. Furthermore, we found that RNA pol II-mediated transcription in the nucleoplasm and in the nucleolus differ greatly in their sensitivity to a transcriptional inhibitor, the latter being blocked at the much lower concentration typical of the one used to specifically inhibit RNA pol I. This suggests that, when transplanted in the nucleolus, RNA pol II might share some of the transcriptional properties of RNA pol I.

In our experiments, the dot-like MS2-GFP signal in the nucleolus forms in large part as a result of transcriptional activity. Indeed, as shown by FRAP curves, the MS2-GFP recovery reaches half of its final value within 4 to 5 minutes, which is comparable to the *t*_1/2_ of 5.5 minutes that was reported in a previous work on the transcriptional dynamics of a CMV-driven MS2-tagged heterologous gene (Yunger et al., 2010). Despite the dynamics that are observed at the transcription site, it should be noted that the average fluorescence recovery never reached 100% at the end of the imaging period (12 minutes after bleaching). Depending on the HeLa transgenic line, and although cell-to-cell variability was high, between 24% and 35% of the pre-bleach MS2-GFP signal failed to re-appear at the transcription spot. This result suggests the existence of a fraction of MS2-GFP-labelled RNAs that are retained at the site of transcription. In agreement with this interpretation, we failed to detect MS2 sequences by RNA FISH either in the cytoplasm or away from the transcription site in the nucleoplasm. We think that the most likely explanation for these observations is that the nucleolar MS2-GFP signal consists of a combination of nascent RNA and RNA that is being processed, degraded or otherwise retained at the site of transcription.

In our imaging data set, the dynamics of RNA pol II activity in the nucleolus are characterized by exponential distributions for both the pulse lengths and the duration of periods of inactivity. Assuming that the rate of processing and/or degradation of the MS2-GFP labelled RNA molecules is constant, the pulsing dynamics are captured by a simple model wherein the genomic loci can be either in a state of inactivity (OFF state) or in one that is permissive of transcription (ON state). In the ON state, transcription fires with a certain probability. We found that nucleolar RNA pol II is active on average for periods of 20.3 minutes. These ON periods are interspersed with periods of inactivity averaging 29.3 minutes, such that the MS2-tagged transcripts are synthesized only ∼40% of the time in our entire imaging data set. It was previously shown that CMV-driven transcription did not occur in pulses (Yunger et al., 2010). The clear pulsing we observed in the transcription of our CMV-driven transgene suggests that transcriptional dynamics might depend as much on the nuclear environment in which transcription takes place as on the regulatory sequences by which it is controlled. In the present case, an obvious cause of pulsing could be the limited availability of RNA pol II in the nucleolus. Indeed, low concentrations of the transcriptional machinery have been linked to the stochasticity of the transcriptional process (Elowitz et al., 2002; Lionnet and Singer, 2012).

Previous work has shown that an epigenetic “memory” of the transcription state of a locus can be inherited through mitosis. In one report, the induction of a transgene was found to occur more rapidly in daughter cells if it had already been induced in the mother cell prior to mitosis (Zhao et al., 2011). The authors suggested that this acceleration is due to the deposition of Lys5-acetylated H4 at the locus and to the subsequent recruitment of BRD4, a protein which recognizes this bookmark. In another instance, the frequency of transcriptional pulsing of a MS2-tagged gene in living *Dictyostelium* cells was found to be more similar between mother and daughter cells than between unrelated cells (Muramoto et al., 2010). Yet another example of mitotic bookmarking is given by the RNA pol I transcriptional machinery itself, which has been found to remain associated with active rDNA during mitosis (Roussel et al., 1996). In view of the marked asynchrony we observed in the post-mitotic re-activation of our transgene in living cells (in ∼80% of the pairs of daughter cells), we believe it unlikely that such mechanisms of epigenetic memory are at work in this case. If they are, then we suggest that they affect differently the two sister chromatids, as we have detected intervals between reappearance of the transcription spot as long as 13.5 hours.

It remains to be determined whether the pulsing of nucleolar RNA pol II transcription we have observed in living cells is a property that applies to RNA pol I-mediated transcription as well. Interestingly, in a previous work using the single molecule RNA FISH technique on fixed *S. cerevisiae* cells, Tan and van Oudernaarden interpreted the distribution of rRNA counts per cell across the population as evidence for “bursty” transcription (Tan and van Oudenaarden, 2010). It is therefore tempting to suggest that our observation of discontinuous transcriptional activity in the nucleolus may reflect cycling in the assembly and disassembly of active chromatin structure in and around the rDNA genes. This cycling might represent an additional regulatory point in the synthesis of ribosomal RNA.

## Acknowledgements

We thank members of our laboratory for helpful discussions. We thank Drs. Robert H. Singer and G. Kondoh for the kind gifts of plasmids. We gratefully acknowledge the financial support of the Czech Science Foundation (GAČR grant P305/12/1246). This work was also supported by Charles University in Prague (PRVOUK P27/LF1/1). V.V.H. was supported by grant 466213 from the Granting Agency of Charles University in Prague (GA UK). The Imaging Center at our institute is supported by the European Regional Development Fund (OPPK CZ.2.16/3.1.00/24010).

## Authors’ contribution

C.L. and V.V.H designed the research, carried out the experiments and wrote the paper. P.K. and G.M.H. implemented the MATLAB-based software for the identification and quantification of transcription spots from 4D image series.

## Legends to Supplementary figures

**Figure S1**. Average intensity profile during FRAP experiments performed on 14 cells of clone MS2^12x^-12. Values (± s.d.) are corrected for total nucleoplasmic signal. Scans were taken every 5 seconds for 14.5 minutes. Maximum intensity projections were used for the evaluation.

**Figure S2**. Automated detection and assessment of transcriptional spots from 4D movies using custom-designed MATLAB-based software. **A)** A representative output image series showing segmented nuclei (circled and numbered in yellow) as well as transcriptional spots (circled and numbered in green). Daughter cells are assigned new identification numbers. Time is indicated in hours and minutes. **B)** Profile of signal strength in the mother (blue) and two daughter cells (red and green). The signal strength is calculated as the signal-to-background ratio based on a Gaussian fit. See text for details.

**Figure S3**. Representation of the imaging data set. ON periods in green, OFF periods in red. Transcriptional traces in the 268 cells that were imaged. Only the interphase portion of the movies are drawn. Time is indicated at the top.

